# The role of ducks in detecting Highly Pathogenic Avian Influenza in small-scale backyard poultry farms

**DOI:** 10.1101/2025.07.23.666292

**Authors:** Steven X Wu, Christopher N Davis, Mark Arnold, Michael J Tildesley

**Affiliations:** The Zeeman Institute, University of Warwick, Coventry, United Kingdom; Mathematics Institute, University of Warwick, Coventry, United Kingdom; School of Life Sciences, University of Warwick, Coventry, United Kingdom; Animal and Plant Health Agency, United Kingdom

## Abstract

Previous research efforts on highly pathogenic H5N1 avian influenza (HPAI) suggest that different avian species exhibit a varied severity of clinical signs after infection. Waterfowl, such as ducks or geese, can be asymptomatic and act as silent carriers of H5N1, making detection harder and increasing the risk of further transmission, potentially leading to significant economic losses. For backyard hobby farmers, passive reporting is a common HPAI detection strategy. We aim to quantify the effectiveness of this strategy by simulating the spread of H5N1 in a mixed-species, small-population backyard flock. Quantities such as detection time and undetected burden of infection in various scenarios are compared. Our results indicate that the presence of ducks can lead to a higher risk of an outbreak and a higher burden of infection. If most ducks within a flock are resistant to H5N1, detection can be significantly delayed. We find that within-flock infection dynamics can heavily depend on the species composition in backyard farms. Ducks, in particular, can pose a higher risk of transmission within a flock or between flocks. Our findings can help inform surveillance and intervention strategies at the flock and local levels.

**Author summary:** We addressed the gap in our understanding of within-flock transmission dynamics of H5N1, particularly for small-scale, backyard farms, where it is reasonably realistic for multiple species of birds to be housed together. These smaller flocks may differ from their larger, industrialised counterparts in their structure and management, and may play a key role in the persistence and spread of H5N1. Notably, we know from the literature that waterfowl, such as ducks, can be asymptomatic after contracting H5N1 and thus act as ‘silent carriers’ of the virus, which could amplify the risk posed to other species of birds, mammals, and humans. We used a stochastic mechanistic model that accounted for such possibilities and simulated the possible outcomes of an outbreak. We found that while the presence of chickens is more likely to lead to high mortality upon infection, ducks can make H5N1 harder to detect within a flock, and thus cause a greater burden of infection, which increases the risk of potential between-flock and between-site transmissions. Our findings are consistent with the current literature and can help inform surveillance and control strategies.

## Introduction

Highly Pathogenic Avian Influenza (HPAI), the H5N1 subtype, remains a threat to global health security as of 2025. The current H5N1 strain (A/goose/Guangdong/1/96) was first isolated from a domestic goose in China in 1996. Since then, numerous large outbreaks occurred in various countries in Asia in 2003, and subsequently led to cases found in Africa, Europe, and the Americas [1, 2]. In 2021, a case of H5N1 was found in a swan sanctuary in Worcester, the United Kingdom, leading to a nationwide outbreak affecting both wild and domestic birds [3]. Although infections are most commonly found among birds, mammals and humans can also be affected. For example, a notable incident occurred in Hong Kong in 1997, where six people died after contracting an infection [2]. In 2024, an outbreak was identified in the US in which dairy cows and two humans contracted H5N1 [4]. These events highlighted the potential of HPAI H5N1 to cause a severe health and economic impact on local or global scales. Therefore, understanding the transmission dynamics of the virus is vital in designing effective control and prevention strategies, and minimising potential human exposure to H5N1, especially in backyard flocks.

HPAI H5N1 is known to have a varied effect depending on the species and breed of the infected bird. Many existing studies, such as Bouma et al. [5] and Yu et al. [6], have shown that species such as chickens and turkeys are likely to exhibit noticeable clinical signs and die within days of an infection. In comparison, there exists some evidence that waterfowl such as ducks and geese can be asymptomatic to H5N1, meaning that no clinical signs would be observed in some cases, while still being highly infectious. This means that ducks can potentially act as ‘silent carriers’ of H5N1, making infections and outbreaks harder to detect promptly [7]. The presence of asymptomatic but infectious birds within a flock may also significantly alter the trajectory of an outbreak. Despite this, relatively few studies have investigated the epidemiological implications of housing multiple bird species together, particularly in small-scale backyard farms where such an arrangement is reasonably realistic [3]. One of the goals of this study is to investigate the potential risk of ‘silent spread’ of H5N1 resulting from the presence of ducks in a small, mixed-species flock, and how the infection dynamics changes for flocks with different species population distributions.

Mathematical and computational models have been widely used to study the spread of infectious diseases, including HPAI H5N1, providing insights into key epidemiological parameters such as the transmission rate and infectious period. In the context of HPAI, several studies have focused on fitting mathematical models to mortality data in large commercial flocks [8, 9], while others have used experimental data to estimate key parameters such as transmission rate, latency period, and infectious period under controlled conditions [10, 11]. A recent review by Kirkeby et al. [12] collected parameter estimates from previous work on HPAI of all subtypes, which were then used for a simulation study [13] to characterise the different epidemiological effects of HPAI subtypes, including H5N1. These works have been instrumental in the design of detection strategies and intervention policies. However, existing current research typically focuses on single-species flocks and overlooks the potentially varied outcomes introduced in mixed-species settings. Furthermore, most current models are geared towards high-density industrialised farms, leaving a gap in our understanding of disease dynamics in small-scale backyard flocks. In this study, we aim to fill this gap by developing a mechanistic model that allows the possibility of inter-species interactions and infection dynamics in mixed-species flocks, enabling exploration of how different population compositions of species influence outbreak probability, detection time, and infection burden.

In this study, we develop a stochastic, mechanistic model to simulate the within-flock spread of HPAI H5N1 in small, mixed-species populations. By varying the proportion of species such as chickens and ducks, we aim to evaluate how species composition influences: (i) the overall disease dynamics assuming no intervention, (ii) the time to HPAI detection via passive surveillance based on mortality signals, and (iii) the total burden of infection, measured by the total time birds spent infectious, up to the point of detection. The potential output of the model is of practical relevance for designing early warning systems, optimising surveillance strategies, and implementing intervention policies. Ultimately, our goal is to improve our understanding of how heterogeneity in species susceptibility, clinical signs, and mortality rate can influence the trajectory and detectability of HPAI outbreaks in non-industrial poultry settings.

## Methods

### Model Construction

We considered a backyard farm that housed either chickens, ducks, or a mixture of both. We used a fixed number of 40 total birds to simulate a smallholder backyard premises. A Susceptible–Exposed–Infectious–Recovered–Dead (SEIRD) model was used as a baseline with added possibilities for a bird to be asymptomatic while infectious.

Therefore, individual birds were divided into the following compartments: susceptible (*S*), exposed (*E*), infectious and symptomatic (*I*), infectious and asymptomatic (*I*^*∗*^), recovered (*R*), and dead (*D*). A subscript was used to denote the species of the bird: *c* for chickens and *d* for ducks (e.g. *S*_*c*_(*t*) denotes the number of susceptible chickens at time *t*). An exposed bird would become infectious at the rate *σ*_*s*_, with the probability of being symptomatic *p*_*s*_, for *s* = *c, d*, where the species was chickens or ducks, respectively. An infectious bird would recover or die with the rate 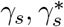, and the probability of death 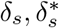, based on the bird species and the existence of clinical signs. The model assumed a homogeneously mixing bird population with a density-dependent contact rate, which is more appropriate for small populations on backyard premises. Therefore, as justified in Fournie et al. [14], the force of infection at time *t* ≥ 0 on species *x* ∈ {*c, d*} was given by:

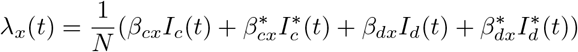

where *N* = 40 was the initial size of the flock, and 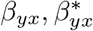 were the transmission rate from symptomatic and asymptomatic birds of species *y* to species *x*, respectively. We assumed all other inter-compartment transitions occur with an exponentially distributed waiting time. Thus, we had the transition rules for chickens:

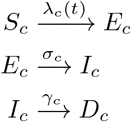

and for ducks:

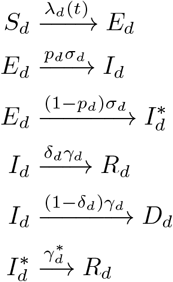

The explanations for the rate parameters are found in Table 1, and the compartmental diagram that captures the transition rules is presented in Figure 1. Note that some transition rules are omitted due to the redundancy caused by the choice of parameters, which can be inferred from Table 1.

**Table 1.** Parameter values used in the model, based on literature sources or specified assumptions.

| Parameter | Description | Value | Source |
| --- | --- | --- | --- |
| $\beta_{cc}$ | Chicken-to-chicken transmission rate | 1.23 (1.11–1.36) | [5, 8] |
| $\sigma_c^{-1}$ | Chicken latency period | 0.24 (0.099–0.48) | [5] |
| $\gamma_c^{-1}$ | Chicken infectious period | 2.1 (1.8–2.3) | [5] |
| $p_c$ | Chicken symptomatic probability | 1 | Assumed [7] |
| $\delta_c$ | Chicken case-fatality probability | 1 | Assumed [7] |
| $\beta_{dd}$ | Symptomatic duck-to-duck transmission rate | 4.1 (2.8–5.8) | [9] |
| $\sigma_d^{-1}$ | Duck latency period | 0.12 (0.03–0.38) | [9] |
| $\gamma_d^{-1}$ | Symptomatic duck infectious period | 4.3 (2.8–5.7) | [9] |
| $p_d$ | Duck symptomatic probability | 0, 0.2, 0.4, 0.6, 0.8 | Sensitivity |
| $\delta_d$ | Symptomatic duck case-fatality probability | 0.70 (0.61–0.78) | [9] |
| $\kappa$ | Rescaling for asymptomatic duck parameters | 2, 3, 4 | Sensitivity |
| $\beta_{dd}^*$ | Asymptomatic duck-to-duck transmission rate | $\frac{\beta_{dd}}{\kappa}$ | Derived |
| $\gamma_d^{*-1}$ | Asymptomatic duck infectious period | $\kappa \gamma_d^{-1}$ | Derived |
| $\delta_d^*$ | Asymptomatic duck case-fatality probability | 0 | Assumed |
| $\beta_{cd}, \beta_{dc}, \beta_{dc}^*$ | Cross-species transmission rates | $\beta_{dd}, \beta_{cc}, \kappa \beta_{cc}$ | Assumed |

Parameter values reflect published estimates where available, with assumptions and derived formulas applied to asymptomatic and cross-species transmission parameters. The rate unit is day<sup>-1</sup>. Sensitivity analysis includes variation in symptomatic probability and rescaling factor $\kappa$ .

**Fig 1.**
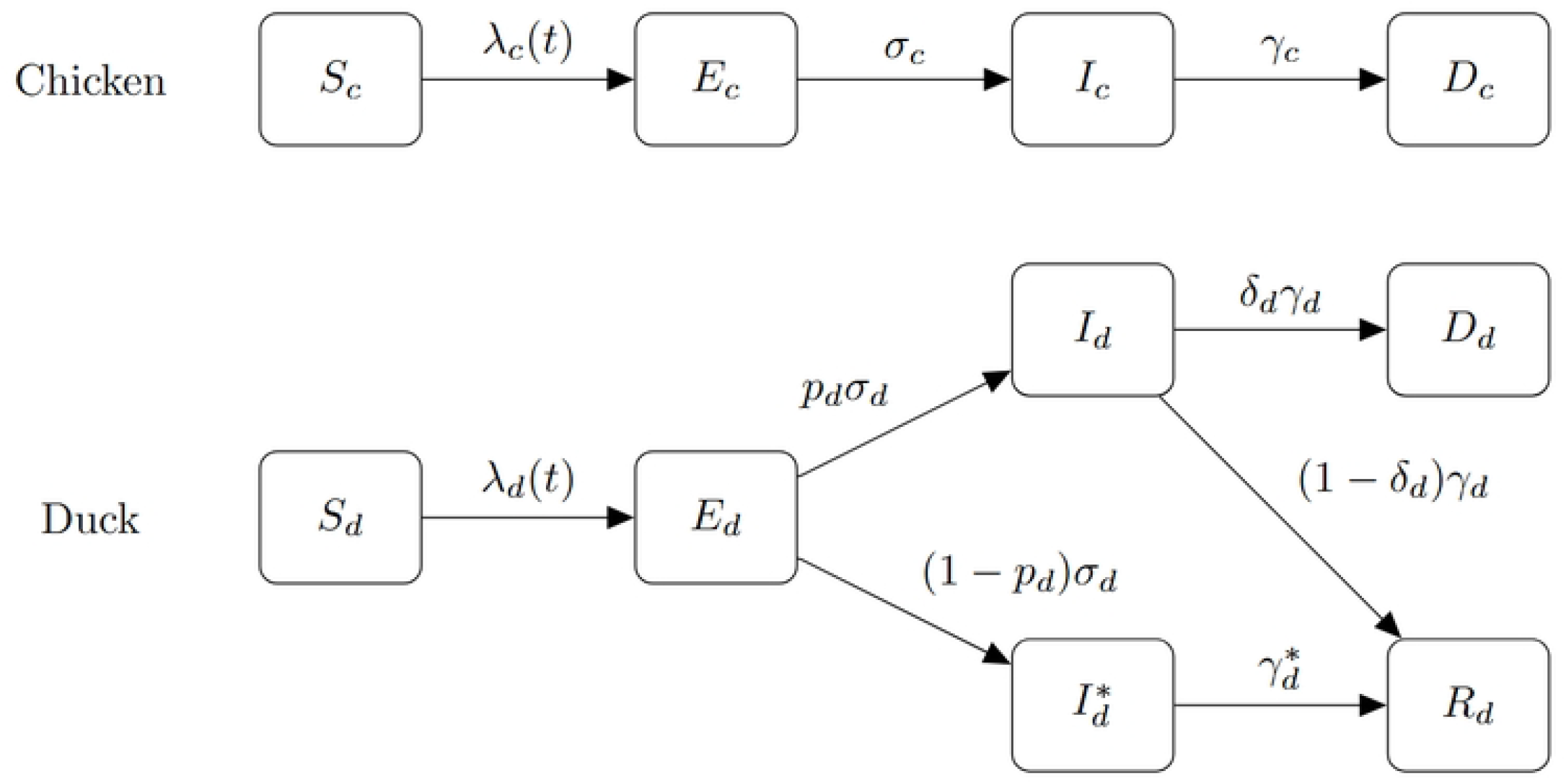
Model Compartmental Diagram. A compartmental diagram of the disease dynamics of a mixed-species flock. The top and bottom parts represent the chicken and duck populations, respectively. We assumed that all infected chickens became symptomatic, while ducks might not exhibit clinical signs with probability (1 − *p*_*d*_).

### Parameter Values

Owing to a lack of real-world data on within-flock infection dynamics for backyard farms, we relied on parameter estimations from the literature to inform our parameter choices. The baseline parameters on chicken infection dynamics (*β*_*cc*_, *σ*_*c*_, *γ*_*c*_) were chosen based on results by Bouma et al. [5] and Tiensin et al. [8]. Specifically, Bouma et al. [5] presented an experimental study that aimed to provide a parameter estimation of an SEIR model for chickens, and Tiensin et al. [8] used Thailand field data to estimate the transmission rate for different values of the infectious period in an SIR model by using the back-calculation method. The values for *σ*_*c*_ and *γ*_*c*_ were taken directly from the experimental study. However, the estimated value for *β*_*cc*_ from the experimental study was based on an inoculated chicken and a susceptible chicken in the same cage, which might not reflect field scenarios. Hence, the value for *β*_*cc*_ was taken from the field study, using the assumed 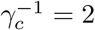, which matched with the experimental study.

Almost all relevant studies suggested that H5N1 is highly lethal to chickens. Experimental results by Bouma et al. [5], Jeong et al. [15], and Forrest et al. [16], as well as reviews such as the work from Kim et al. [7], implied a 100% mortality rate for infected chickens. While works from Spekreijse et al. [11] and Yu et al. [6] demonstrated that a small proportion of chickens might be asymptomatic depending on the strains of H5N1, but near 100% mortality rate for chickens with clinical signs was still observed. Therefore, we assumed in our model that chickens would always show clinical signs and eventually die of H5N1. For ducks, it was found that the possibility of showing clinical signs and dying from H5N1 varies depending on the breed of the duck [7, 17], the strains of the virus [6, 17–19], or the route of infection [15]. Therefore, we conducted a sensitivity analysis on the duck symptomatic probability (*p*_*d*_) to account for the different scenarios, choosing values *p*_*d*_ = 0, 0.2, 0.4, 0.6, and 0.8.

Current literature on parameter estimation of infection dynamics in duck flocks is scarce. One study on HPAI H5N8 from Vergne et al. [9] estimated using field data that the transmission rate was 4.1, the latency period was 0.12, the infectious period was 4.3, and the case-fatality rate was 0.7. Although the HPAI serotype was different, the results from the study were reasonably in agreement with an experimental study from van der Goot et al. [10]. The estimated *R*_0_ number for duck flocks was also significantly larger than that of chicken flocks, in agreement with a recent review by Kirkeby et al. [12]. Given the high case-fatality rate assumed in the study, we assumed that the viral strain of focus in the work from Vergne et al. [9] was highly pathogenic to ducks, so the parameter estimations provided would apply specifically to symptomatic ducks.

For asymptomatic ducks, there was a lack of parameter estimation for the transmission rate or infectious period. Based on the work from Hulse-Post et al. [18], we expected that the infectious period for asymptomatic ducks could be as long as 17 days. Therefore, we defined a rescaling factor *κ* = 3, where we assumed that the transmission rate 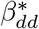 and infectious period 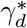 from asymptomatic ducks were 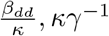 respectively, which ensured the value for *R*_0_ was fixed. While *κ* = 3 will be assumed for all simulations and results for the rest of the paper, we also tested cases when *κ* = 2 and 4 for sensitivity analysis (See Figures S5 Fig-S8 Fig from supporting information). Finally, given the homogeneous-mixing population assumption, we set the cross-species transmission rate *β*_*cd*_ = *β*_*dd*_, *β*_*dc*_ = *β*_*cc*_, and 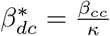, assuming the same contact rate between any two birds and the probability of infection dependent on the receiving bird.

### Simulations and Analysis

The model was simulated using the Gillespie algorithm [20]. Given a set of parameter choices, we set the number of chickens in the population to be 0, 10, 20, 30, and 40, out of 40 total birds, with the remainder being ducks. For the initial condition, one bird was chosen at random to become exposed, and the rest of the flock remained susceptible. The simulation then continued until no disease was present in the system. We performed 5000 realisations of the simulation model and stored them as a time series for each set of parameter choices and species compositions, which can be directly analysed by extracting summary statistics.

Among the 5000 simulations, we considered those that resulted in five or more birds ever being infected as an outbreak. We then filtered out the time series with no outbreaks and considered the interquartile range of the detection time: the day when the farm owner detects a certain number of deaths in the flock. We emulated the behaviours of different types of farm owners based on the level of cautiousness and likelihood of compliance, by considering scenarios where they would report a detection when two, four, or six birds died in the flock. The detection time for each detection rule was then recorded. We also calculated the undetected burden of infection within the flock, defined by the total time birds spent infectious up to the time of detection, measured in bird-days. This was done for every simulation with the death threshold for detection set at two, four, or six, similar to the detection time calculation.

## Results

### Summary Statistics

We investigated the final size of the simulations, recording the final death number (Figure 2 and 3 for each simulation. In most cases, most birds in the flock would be infected at some point during a successful outbreak (meaning that they will be either dead or recovered in the end). A higher duck symptomatic probability tends to lead to a larger number of deaths in a flock. This is evident from both the histograms in Figure 2 and 3 and the heat map in Figure 4. Interestingly, Figure 4 suggests that flocks with 30 chickens and 10 ducks result in the highest average number of deaths for all values of duck symptomatic probability.

**Fig 2.**
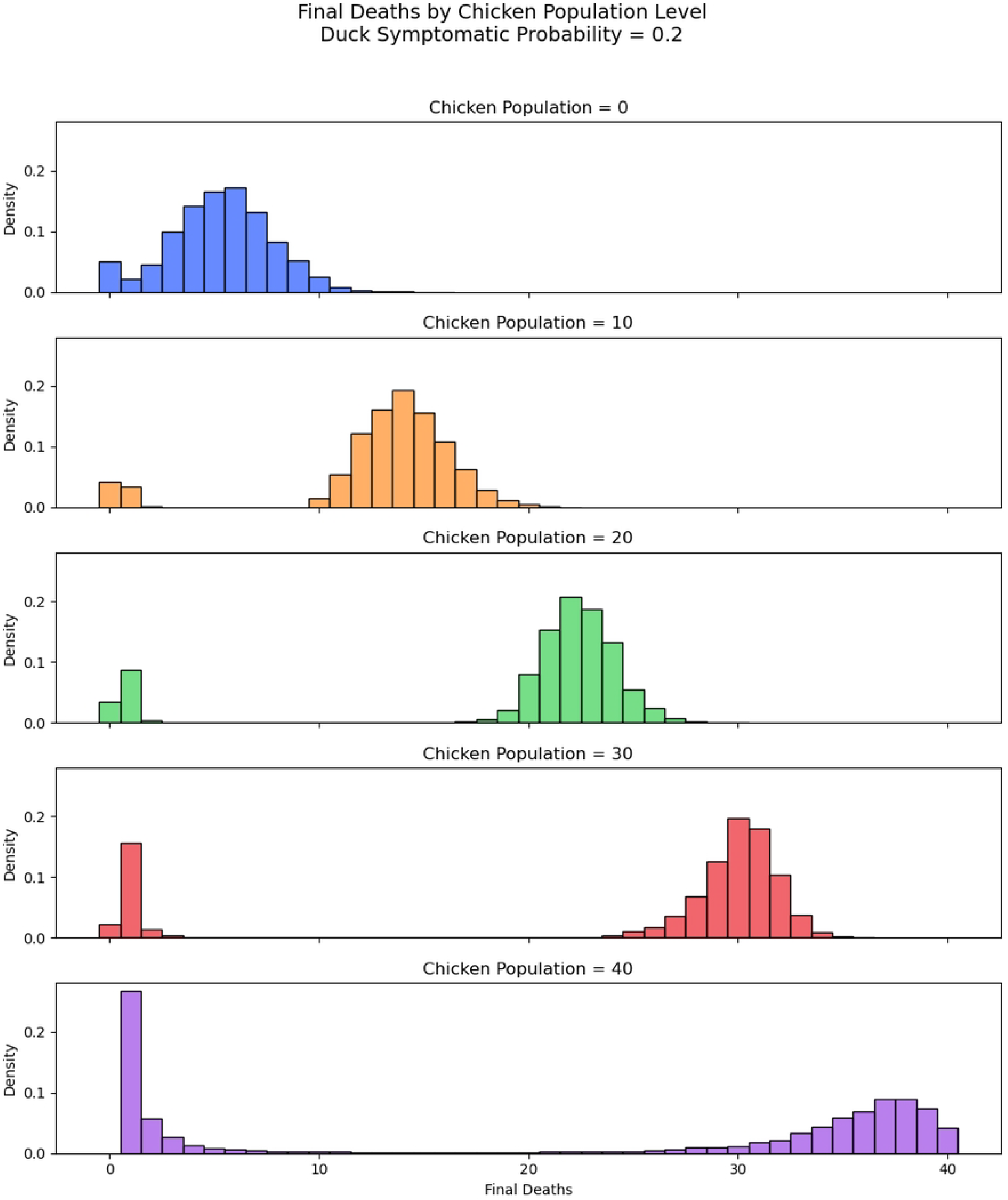
Histogram Final Deaths. *p*_*d*_ = 0.2 Histogram of distribution of final death numbers with varied number of chickens in flocks of size 40. Duck symptomatic probability set to 0.2.

**Fig 3.**
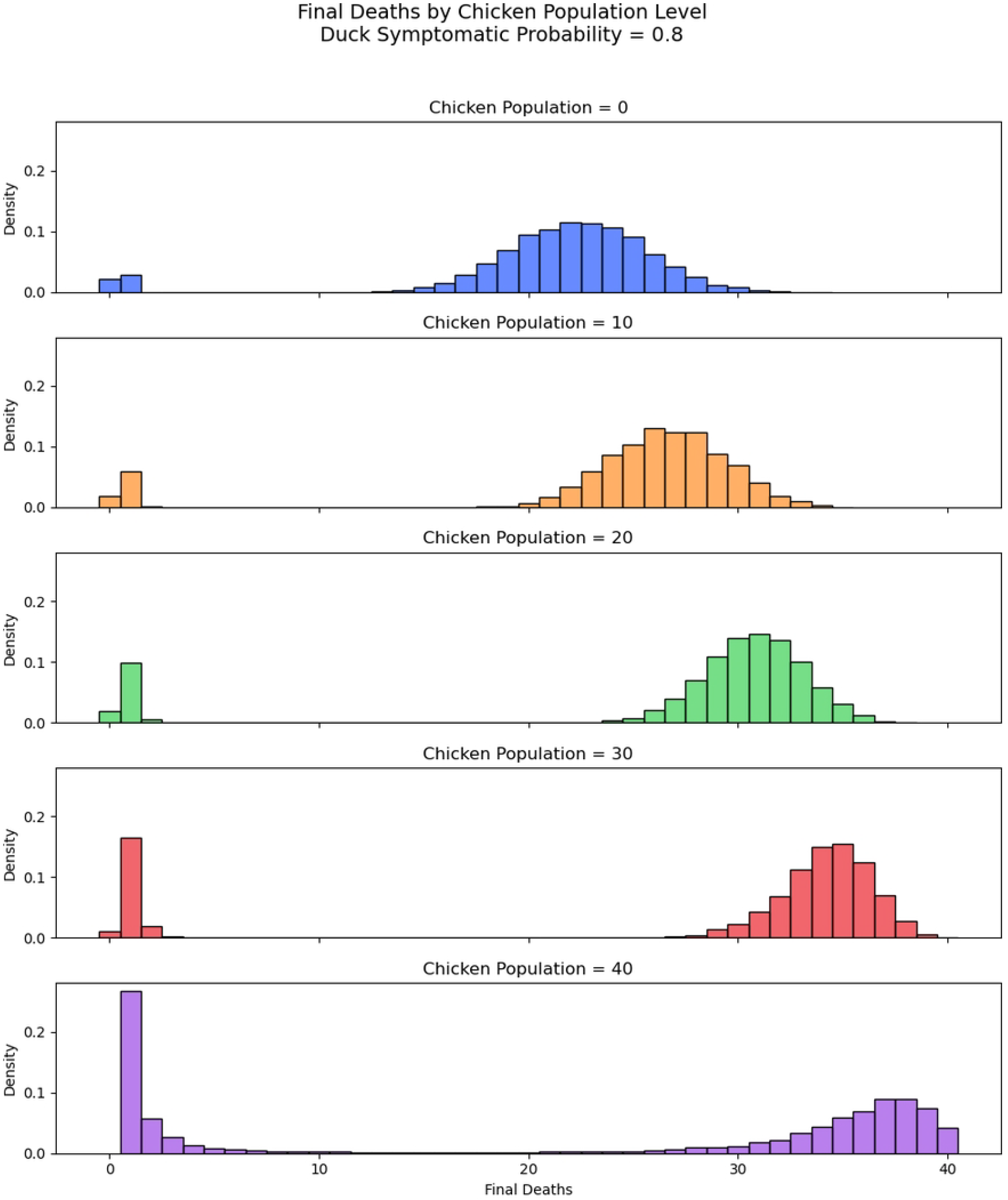
Histogram Final Deaths. *p*_*d*_ = 0.8 Histogram of distribution of final death numbers with varied number of chickens in flocks of size 40. Duck symptomatic probability set to 0.8.

**Fig 4.**
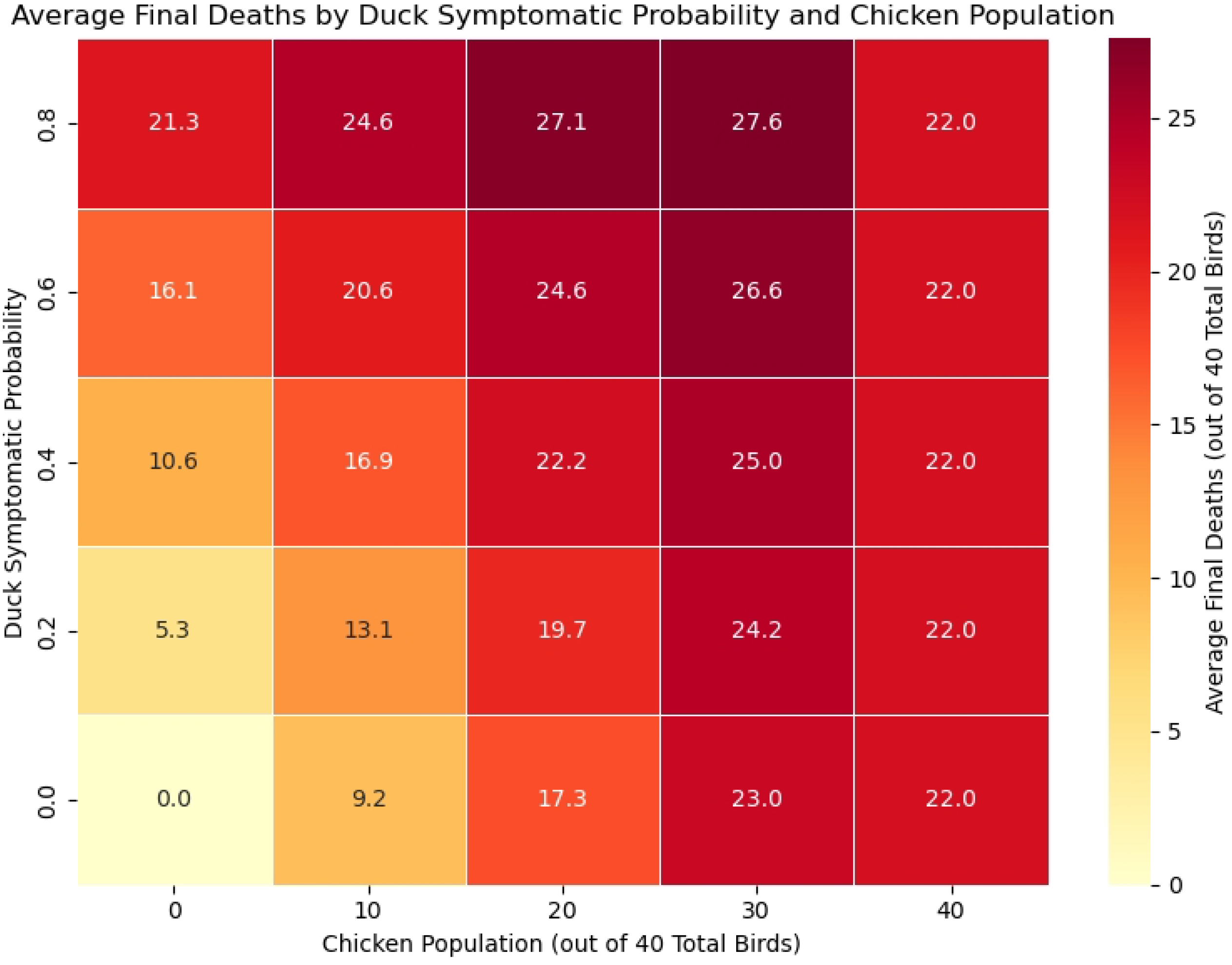
Heat Map Final Deaths. Heat map showing the average number of deaths in all simulation results given the parameter combinations of chicken populations and duck asymptomatic probability.

### Detection Time

We found that the outbreak detection time was highly dependent on the choice of parameters in the sensitivity analysis, specifically the probability of a duck showing clinical signs upon infection. Firstly, as per expectations, farmers who are able to detect H5N1 with lower death thresholds had an overall shorter detection time since the initial infection, as shown in Figure 5. We found that the difference between detection thresholds in duck-only flocks is most significant if the duck’s symptomatic probability is small (Figure 6 and top plot of Figure 5). We also found that duck-only flocks can result in overall longer detection times, with H5N1 left undetected for 10 days (median), and for as long as 40+ days in the worst-case scenarios. In comparison, the detection time is lower if chickens are present in the flock. Notably, detection is expected within approximately 7 days since the initial infection for chicken-only flocks. However, if the duck symptomatic probability is high, due to the higher mortality rate for ducks, the detection time is significantly lowered (bottom plot of Figure 5). The presence of chickens instead increases the detection time slightly, but the overall expected detection time is still less than approximately 7 days.

**Fig 5.**
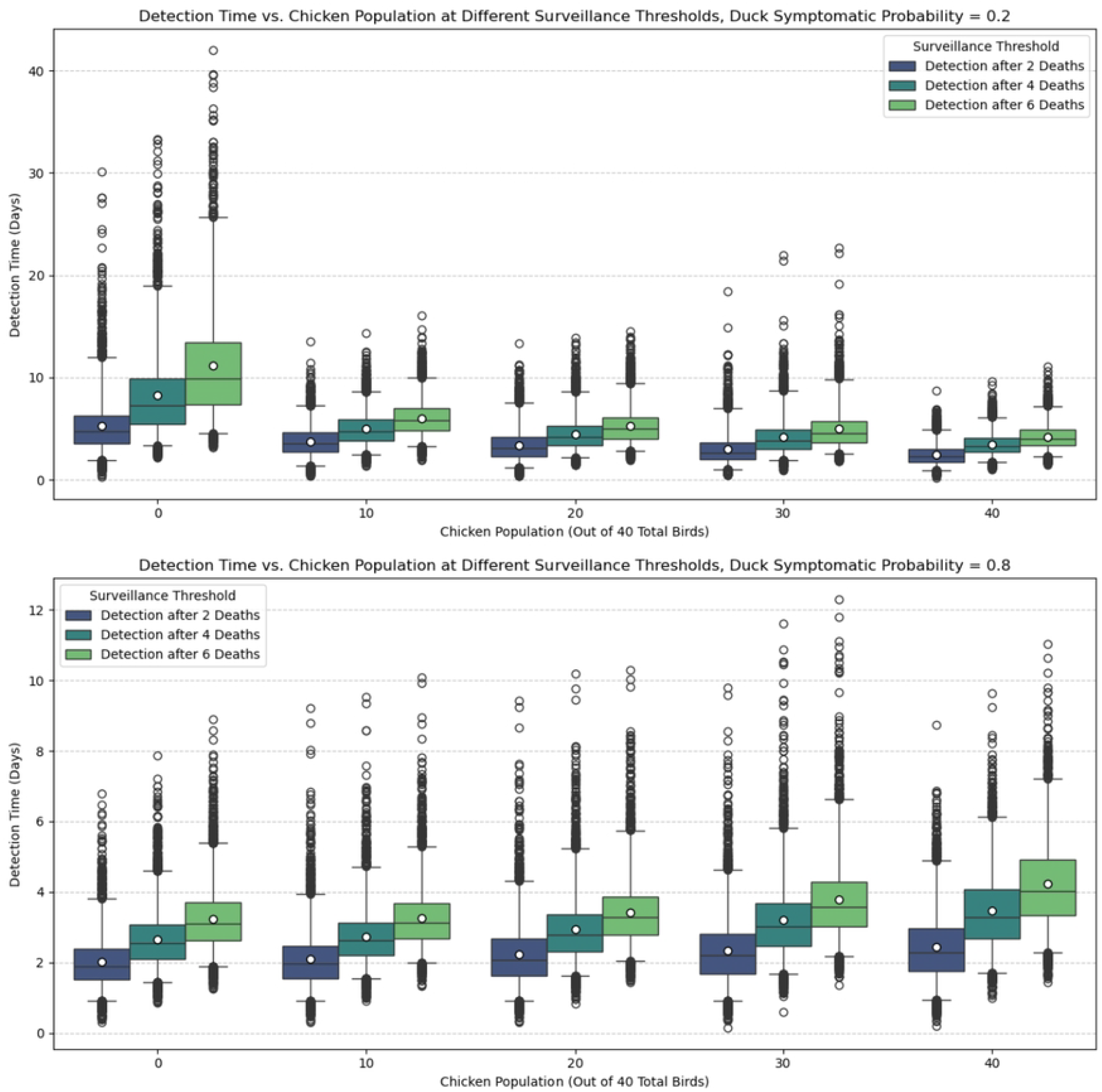
Box Plot Detection Time. Detection time for different death thresholds for detection and chicken population out of 40 total birds. Top plot: the duck’s symptomatic probability *p*_*d*_ = 0.2; Bottom plot: the duck’s symptomatic probability *p*_*d*_ = 0.8. The boxes indicate the interquartile range (25th and 75th percentiles), and the whiskers indicate the 2.5th and 97.5th percentiles. The white dot in the box indicates the average detection time, which also corresponds to the fourth and first row of Figure 6 (for detection after 2 deaths).

**Fig 6.**
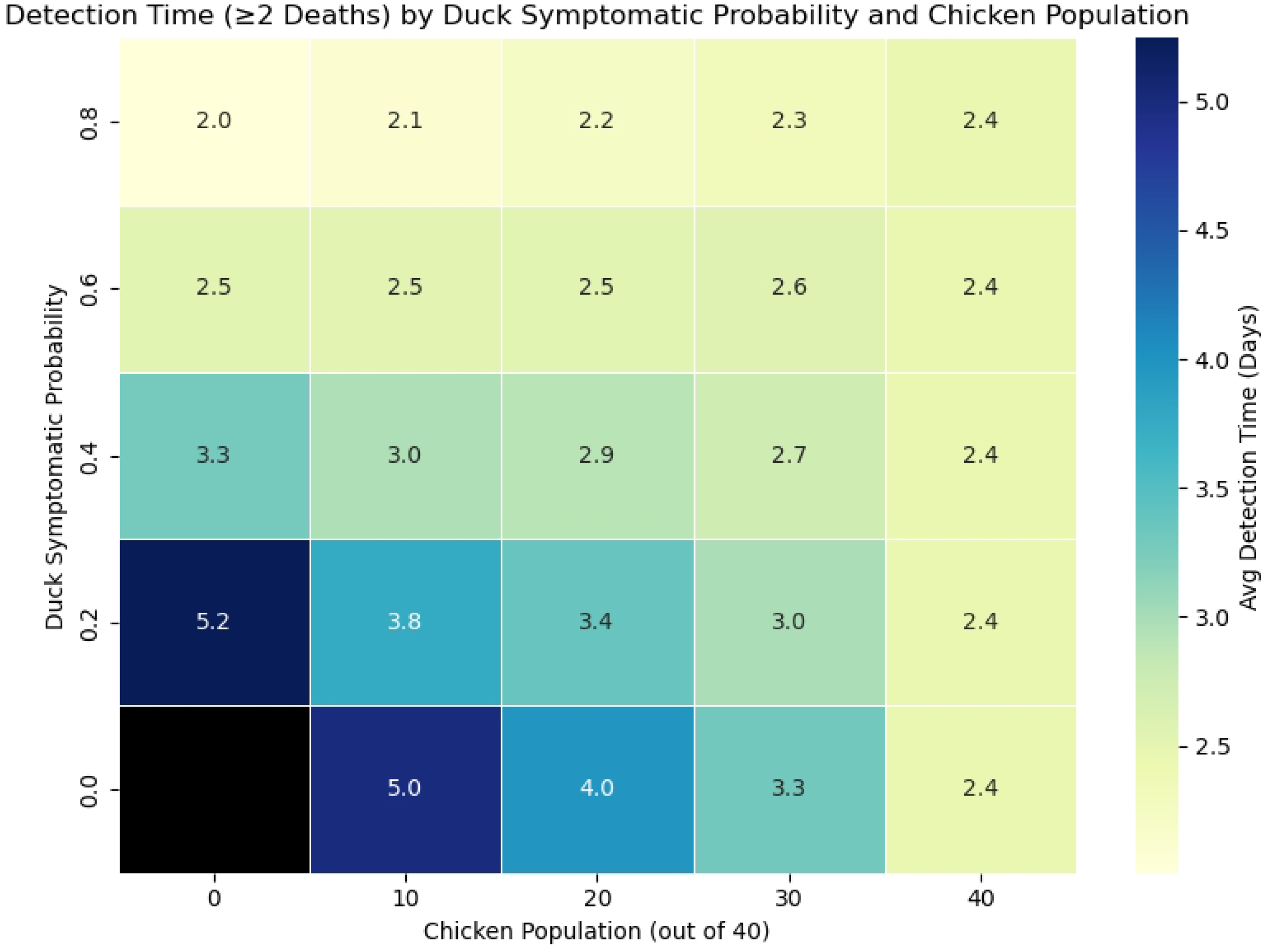
Heat Map Detection Time. Heat map showing the average detection time (with detection occurring when two deaths have occurred) in all simulation results given the parameter combinations of chicken populations and duck symptomatic probability. Note that no detection would occur for duck flocks with duck symptomatic probability equal to zero (Black grid).

### Burden of Infection

In general, due to the long infectious period, the transmissibility of H5N1, and the potentially delayed detection time in duck flocks, their presence in a flock contributes more to the burden of infection compared to chickens. If the duck symptomatic probability is low, the total time birds spend infectious is significantly higher in duck-only flocks, approximately 18 times higher than that of chicken-only flocks (top plot of Figure 7). While the number is reduced in the case of high duck symptomatic probability, a duck-only flock would still have approximately 4 times the burden of infection than chicken-only flocks (bottom plot of Figure 7). Note that we only accounted for time series with detected outbreaks, so if an outbreak occurred but detection failed, then the burden of infection accounting for such cases will be higher in duck-only flocks. As before, the heat map for the burden of infection before detection is shown in Figure 8. The result is consistent with the findings in Figure 7 for other sets of parameter choices.

**Fig 7.**
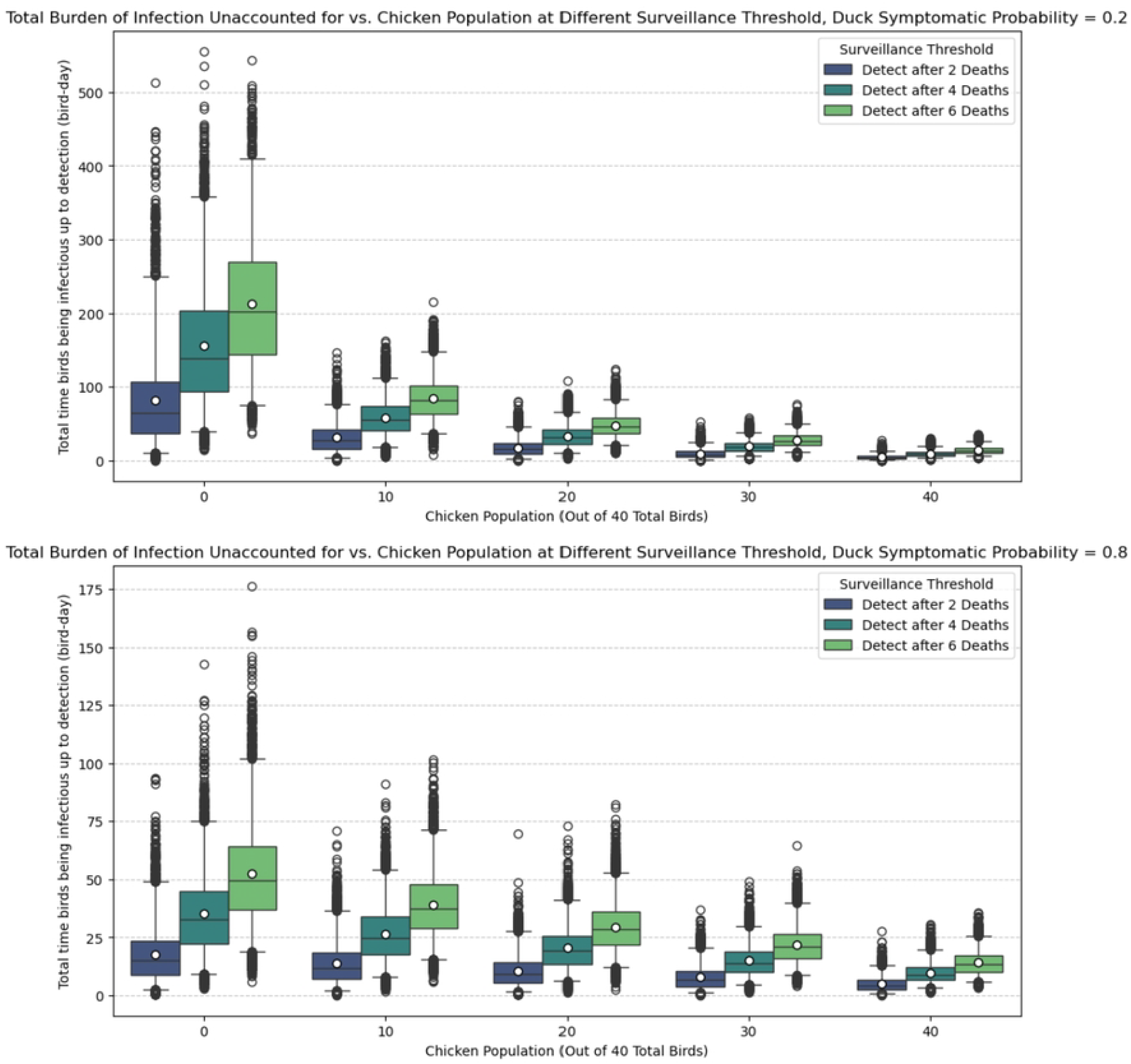
Box Plot Burden of Infection. Undetected burden of infection (total time birds spent infectious up to the point of detection) for different death thresholds for detection and chicken population out of 40 total birds. Top plot: the duck’s symptomatic probability *p*_*d*_ = 0.2; Bottom plot: the duck’s symptomatic probability *p*_*d*_ = 0.8. The boxes indicate the interquartile range (25th and 75th percentiles), and the whiskers indicate the 2.5th and 97.5th percentiles. The white dot in the box indicates the average detection time, which also corresponds to the fourth and first row of Figure 8 (for detection after 2 deaths).

## Discussion

Our study indicates that ducks, or other similar species, may pose a substantial risk in the context of within-flock infections of HPAI. In particular, duck-only flocks have a higher chance of having an H5N1 outbreak (see Figure S3 Fig and S4 Fig from supporting information), causing delayed detection (Figure 5 and 6) and increasing the risk of undetected outbreaks, and having a significantly higher burden of infection up to the point of detection (Figure 7 and 8), especially if ducks are more likely to be asymptomatic and thus can act as a ‘hidden’ source of infection. Additionally, having a small population of ducks in a chicken-dominant flock can amplify mortality, as shown in Figure 4, where maximum final deaths are observed for flocks with 30 chickens and 10 ducks given a fixed duck symptomatic probability. We hypothesise that ducks can act as a more persistent intermediate source of infection, which could lead to more widespread infections when a larger chicken population coexists with the ducks. In general, our findings are consistent with many other studies, in which ducks are shown or implied to be a more significant risk factor for HPAI H5N1 transmission [7, 9, 21].

**Fig 8.**
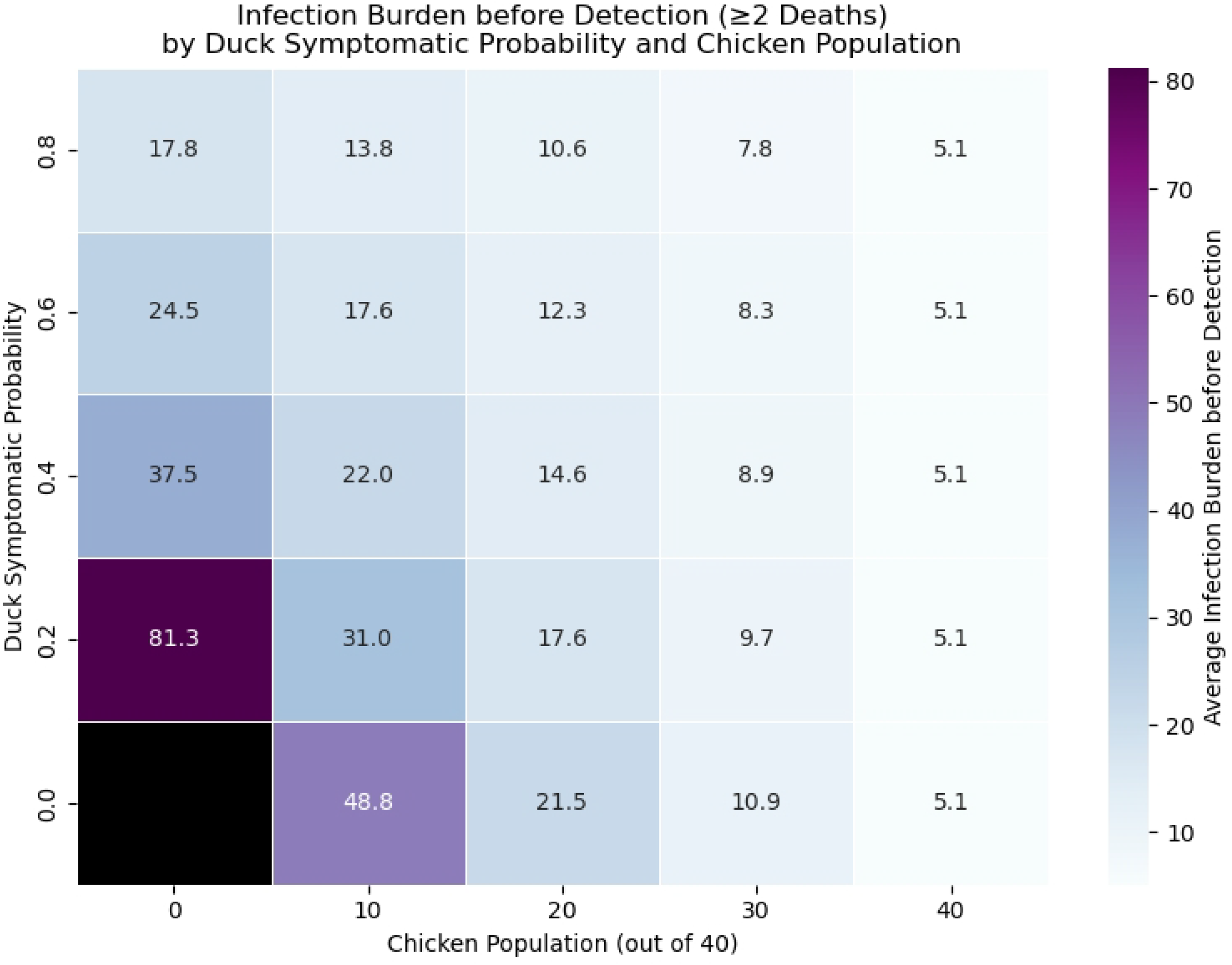
Heat Map Burden of Infection. Heat map showing the average undetected burden of infections (with detection occurring when two deaths have occurred) in all simulation results given the parameter combinations of chicken populations and duck asymptomatic probability. Note that no detection would occur for duck flocks with duck symptomatic probability equal to zero (Black grid).

The most sensitive parameter in our study is *p*_*d*_, the probability that a duck will show clinical signs after infection. Specifically, detection evasion tends to only occur in duck-only flocks if *p*_*d*_ is small. This is likely due to low duck mortality, and most birds in the flock will recover without being detected. For larger values of *p*_*d*_, we found that the detection time becomes shorter in duck-dominant flocks than in chicken-dominant flocks. A possible explanation is that ducks have a higher transmission rate and a longer infectious period, increasing the rate of death among birds. We also saw a lower infection burden when we increased the value of *p*_*d*_, which is potentially due to the reduced detection time.

One of our main assumptions regarding detection time is the criteria for detection. We assumed that a passive surveillance approach, where farmers voluntarily report a suspected infection in their flocks, has been implemented. We had chosen this approach as opposed to other strategies recommended for larger commercial flocks [21], because we believe that backyard farmers are more likely to have direct contact with the birds and therefore can observe signs and/or death reasonably efficiently. However, farmers’ knowledge and attitudes towards HPAI can vary significantly and are difficult to quantify. A recent survey study by McClaughlin et al. [3] highlighted the lack of knowledge of the clinical signs of HPAI among UK birdkeepers. Hence, we have chosen the detection criterion to be ‘a certain number of deaths found in the flock’. We chose the threshold to be two, four, or six deaths to represent a likely range of farmer behaviour, but a highly knowledgeable bird owner may be able to identify clinical signs in birds before their deaths, and a less knowledgeable farmer may only report when there is a large number of deaths. To address the uncertainties of detection criteria, more surveys and investigations are required to better quantify farmers’ behaviours, as has been implemented in other modelling studies [22].

Our choice of simulation model differs from some other studies on commercial farms [9, 21] as we have excluded natural deaths of birds in our model. Unlike in a commercial farm, where the process of raising birds can be highly automated, backyard hobby farmers tend to know and have close contact with their flocks. Hence, while natural and H5N1-induced mortality can be difficult to differentiate in a large-scale farm, we assumed that small-scale bird keepers can identify an ‘unnatural’ death with relative ease, so the existence of natural mortality in our model makes little impact.

A limitation of our study is the choice of parameters in our model. Many parameter choices for our simulation were taken from either experimental studies (e.g. Bouma et al. [5]) or estimations from field data (e.g. Tiensin et al. [8], Vergne et al. [9]). Such parameters may change depending on the biosecurity measures and housing conditions of each individual flock. Notably, the inter-species transmission rate may be lower than what we assumed if chickens and ducks are kept separately, which invalidates our assumption of a homogeneously mixing population of birds. Some parameters, such as the probability of chickens showing clinical signs and dying from H5N1, were based on evidence from the literature and may be challenged as well.

From a policy-making perspective, our findings suggest that species composition within flocks has important implications for HPAI control strategies. Given the higher transmission potential associated with ducks, or waterfowl in general, we hypothesise that strategies such as active surveillance should prioritise duck-dominant flocks if the objective is to minimise the number of affected farms. However, if the goal is to reduce overall mortality, particularly when vaccination is a considered option, then prioritising chicken-dominant flocks, especially those also in close contact with ducks, may be more effective, as chickens tend to die of H5N1 at a faster rate and probability.

In conclusion, this study presents an exploratory analysis of how species composition may influence the within-flock transmission dynamics of HPAI. Our finding suggests that ducks, or waterfowl in general, are likely to play a more significant role in transmitting HPAI, while chickens or other similar species tend to have faster and more frequent mortalities, which is consistent with the current literature. Future work on parameter estimation of infection dynamics between various species of birds can be useful for further investigation. Another potential direction of research is to consider alternative surveillance and control strategies by taking the different transmissibility of H5N1 from various bird species into account.

## Supporting information

**S1 Fig.**
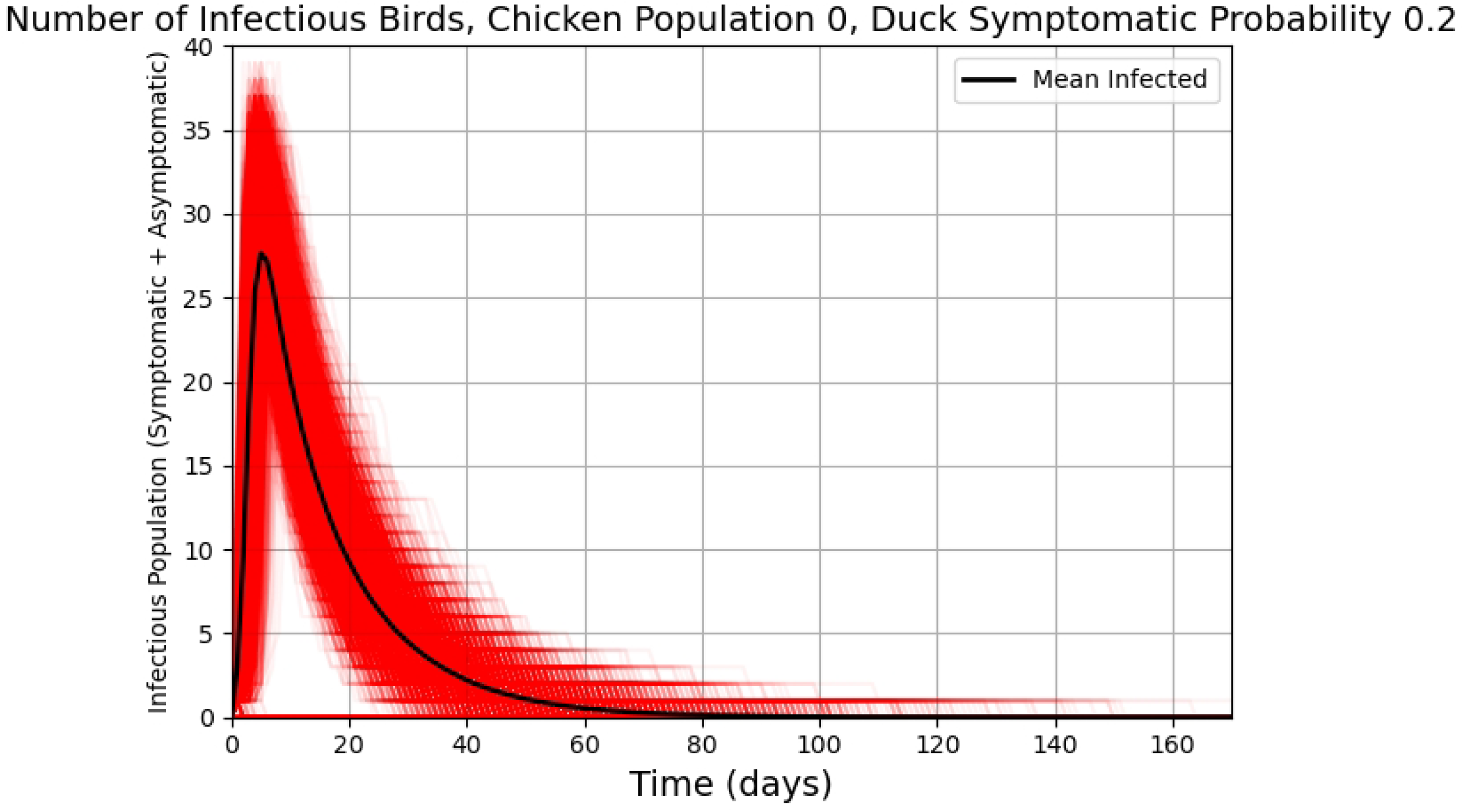
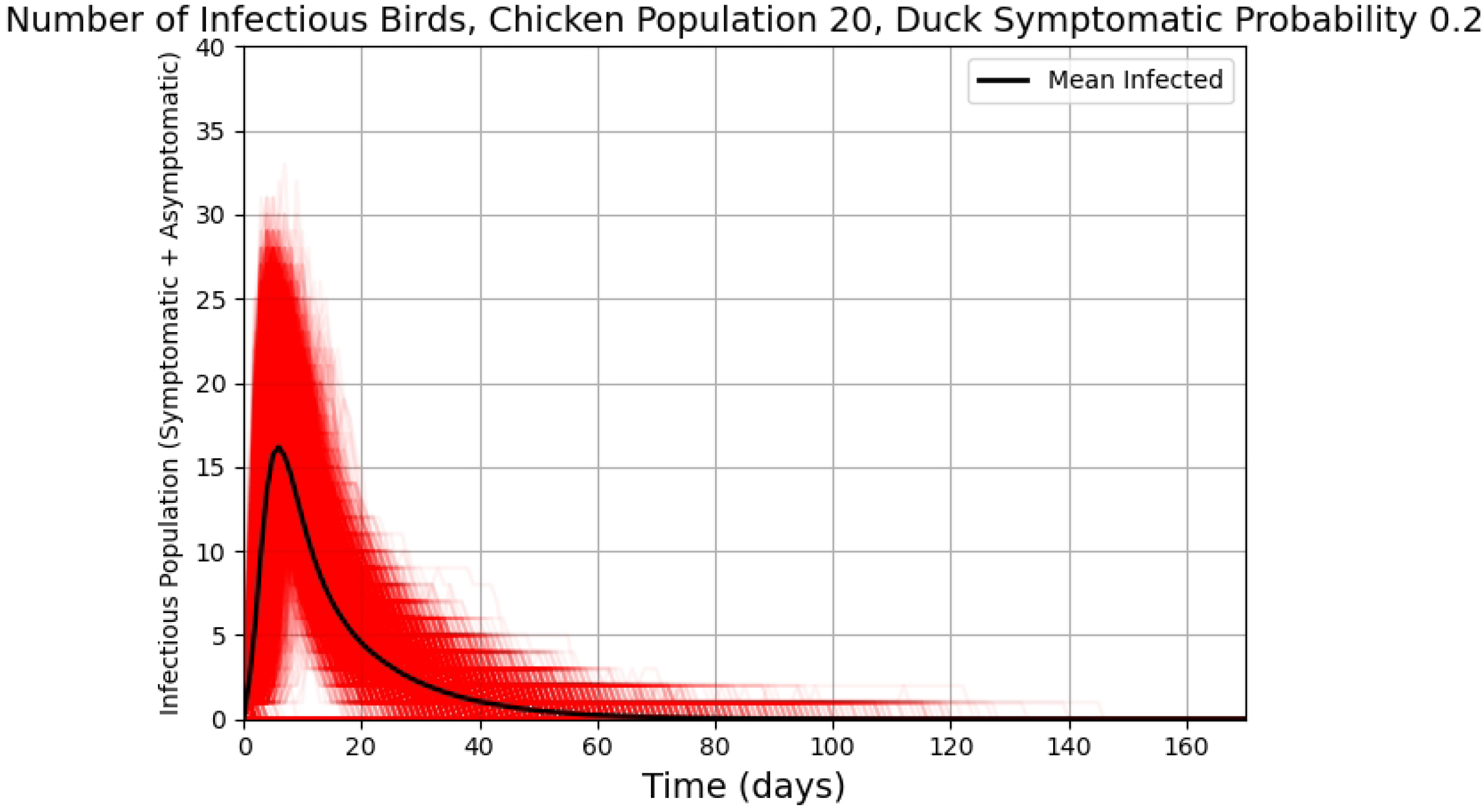
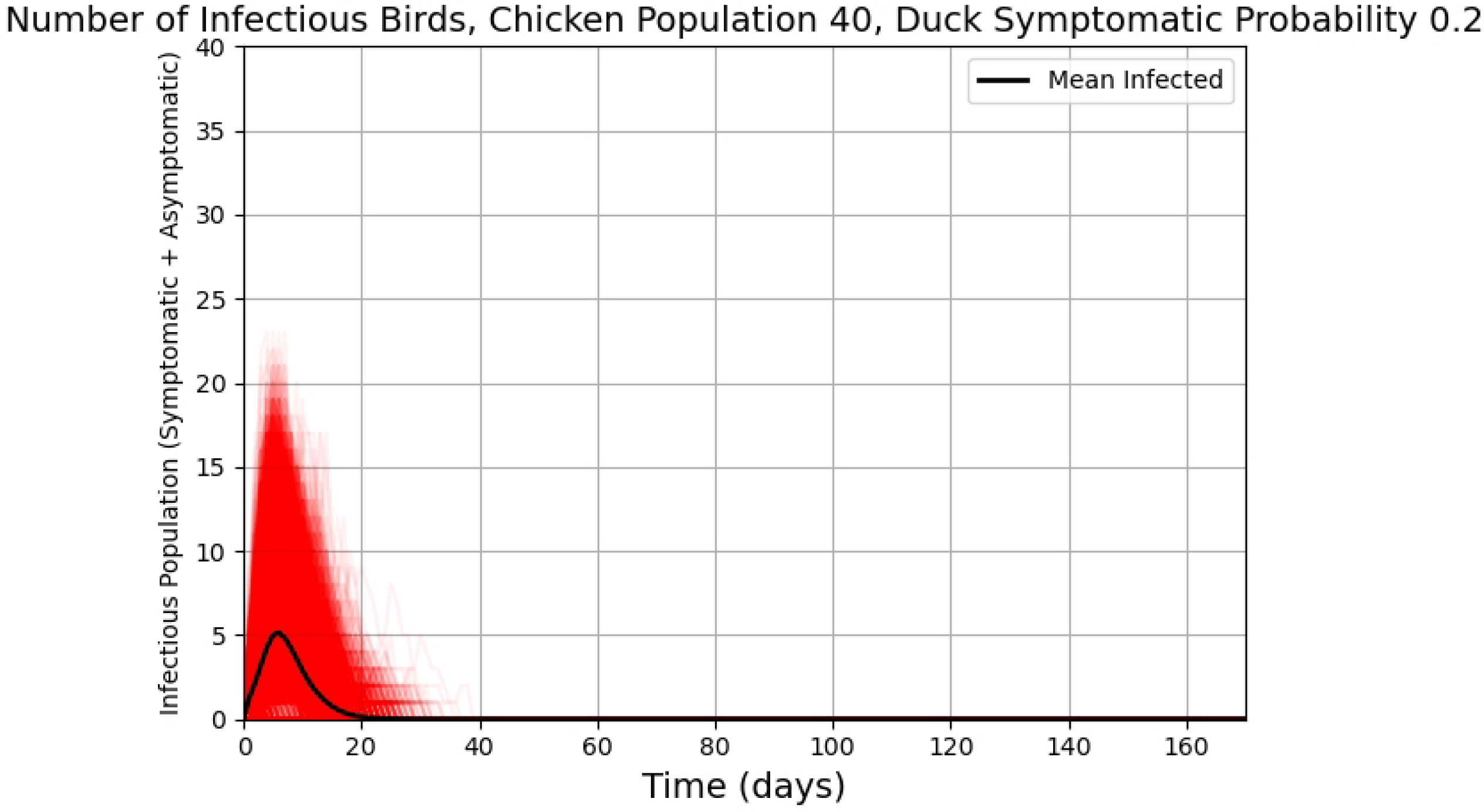
Infectious Birds Time Series. Number of infectious birds over time, for fixed *p*_*d*_ = 0.2. Red trajectories shows the time series directly, and the black trajectory is plotted by calculating the average across all time series on each day. Top: all 40 birds are ducks, middle: 20 chickens and 20 ducks, bottom: all 40 birds are chickens.

**S2 Fig.**
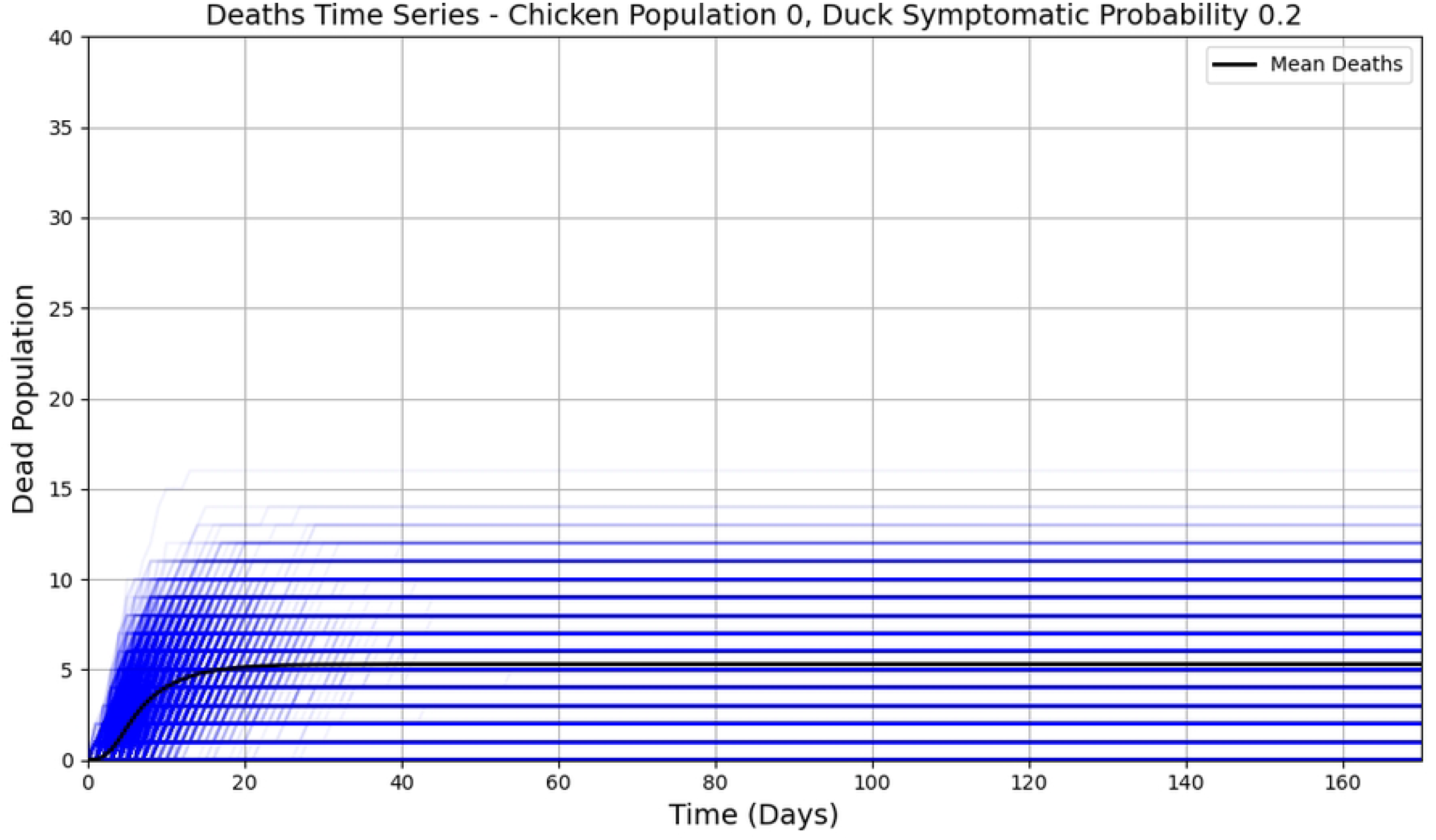
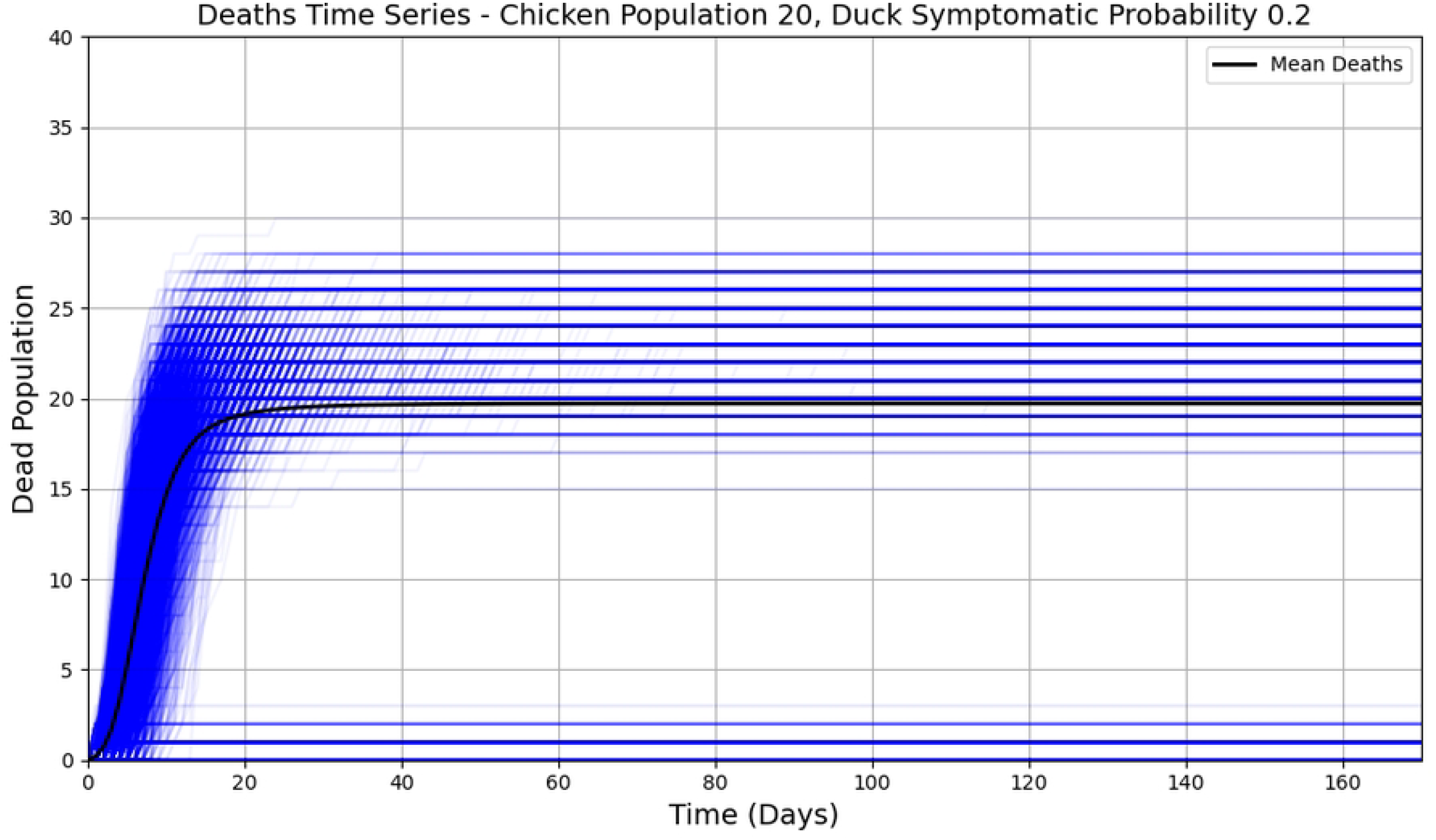
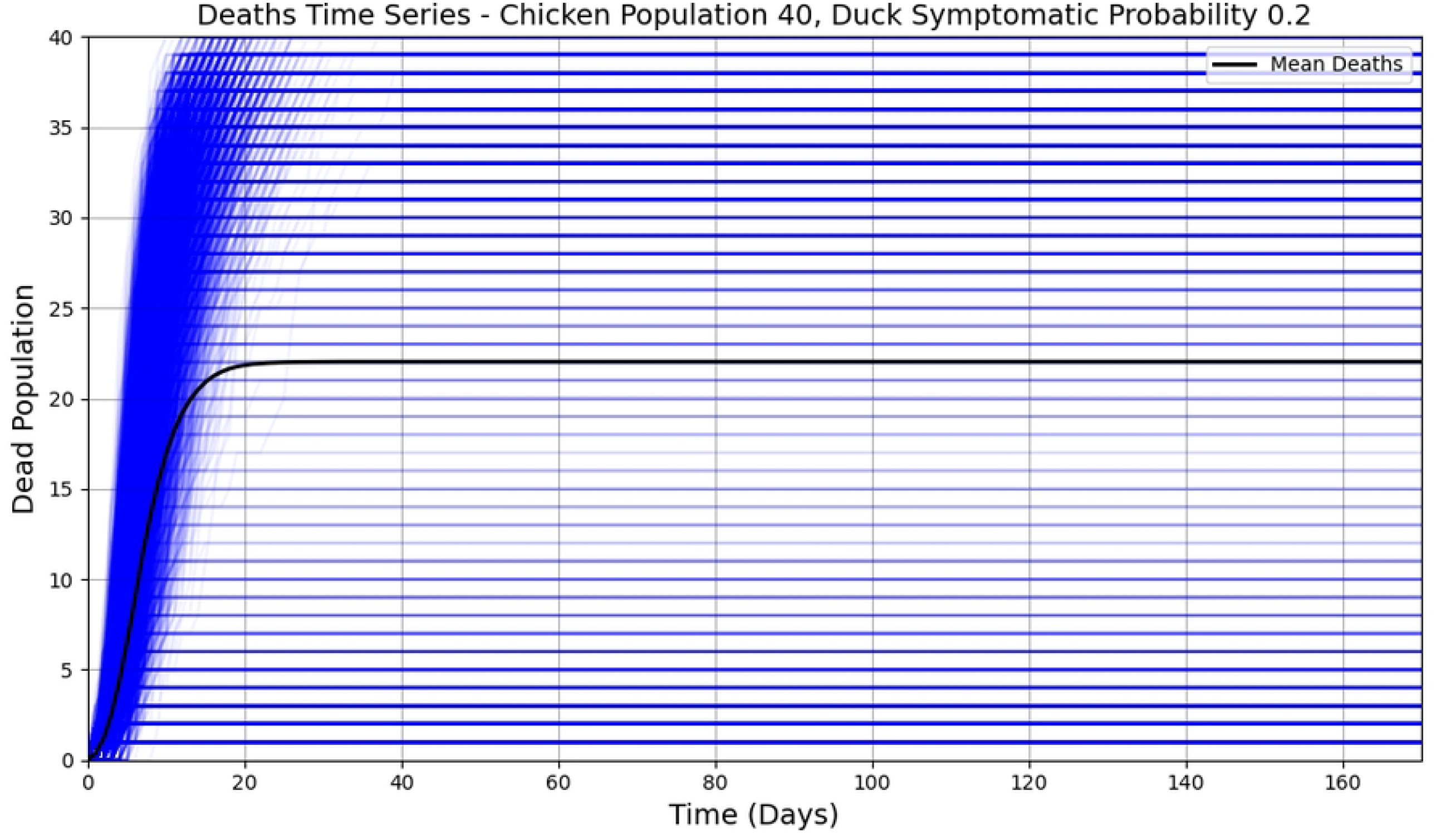
Dead Birds Time Series. Number of dead birds over time, for fixed *p*_*d*_ = 0.2. Blue trajectories shows the time series directly, and the black trajectory is plotted by calculating the average across all time series on each day. Top: all 40 birds are ducks, middle: 20 chickens and 20 ducks, bottom: all 40 birds are chickens.

**S3 Fig.**
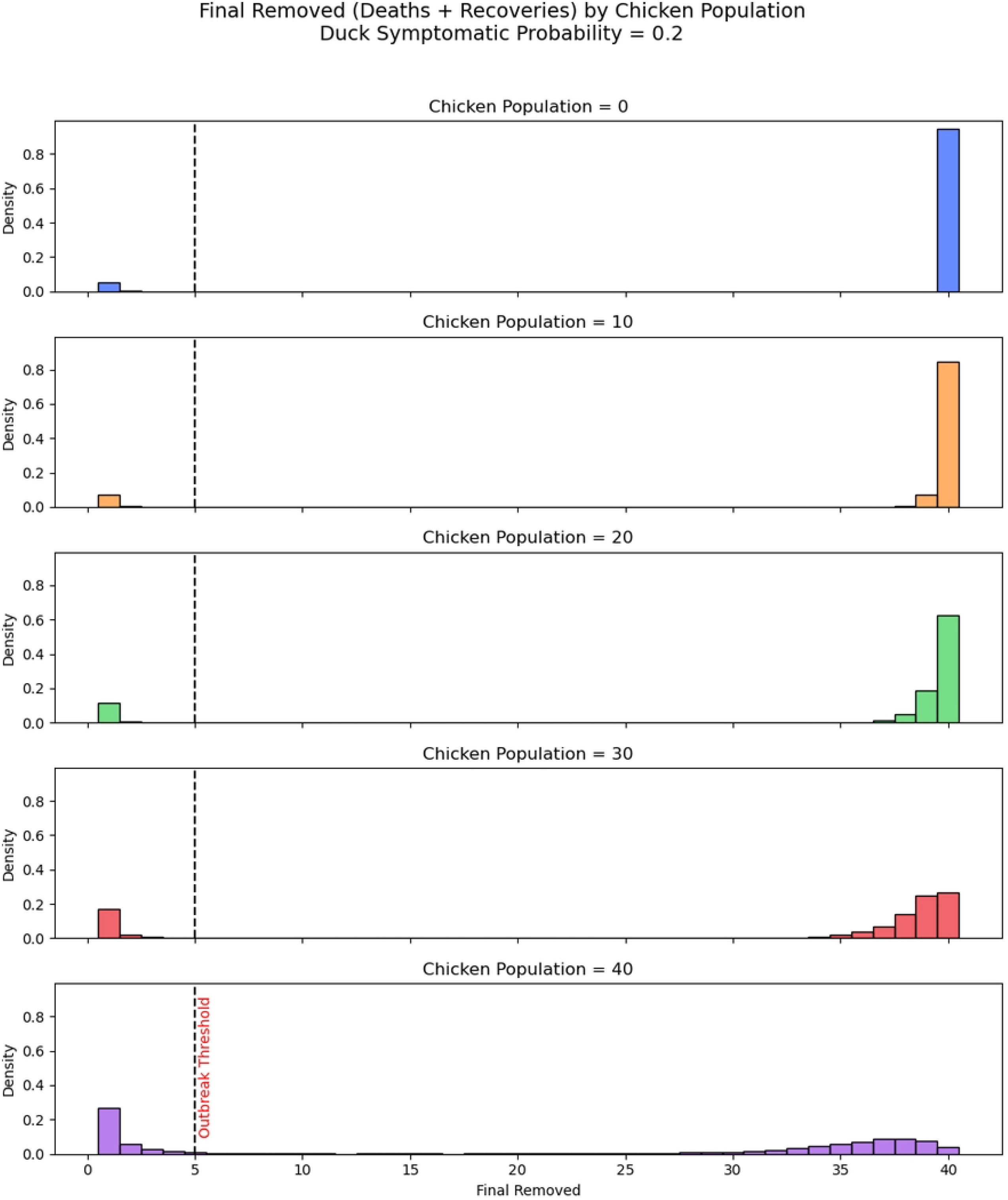
Histogram Final. *p*_*d*_ = 0.2 Histogram of distribution of final removed numbers (dead or recovered) with varied number of chickens in flocks of size 40. Duck symptomatic probability set to 0.2. The vertical dashed line indicates time series that are considered an outbreak (To the right of the dashed line).

**S4 Fig.**
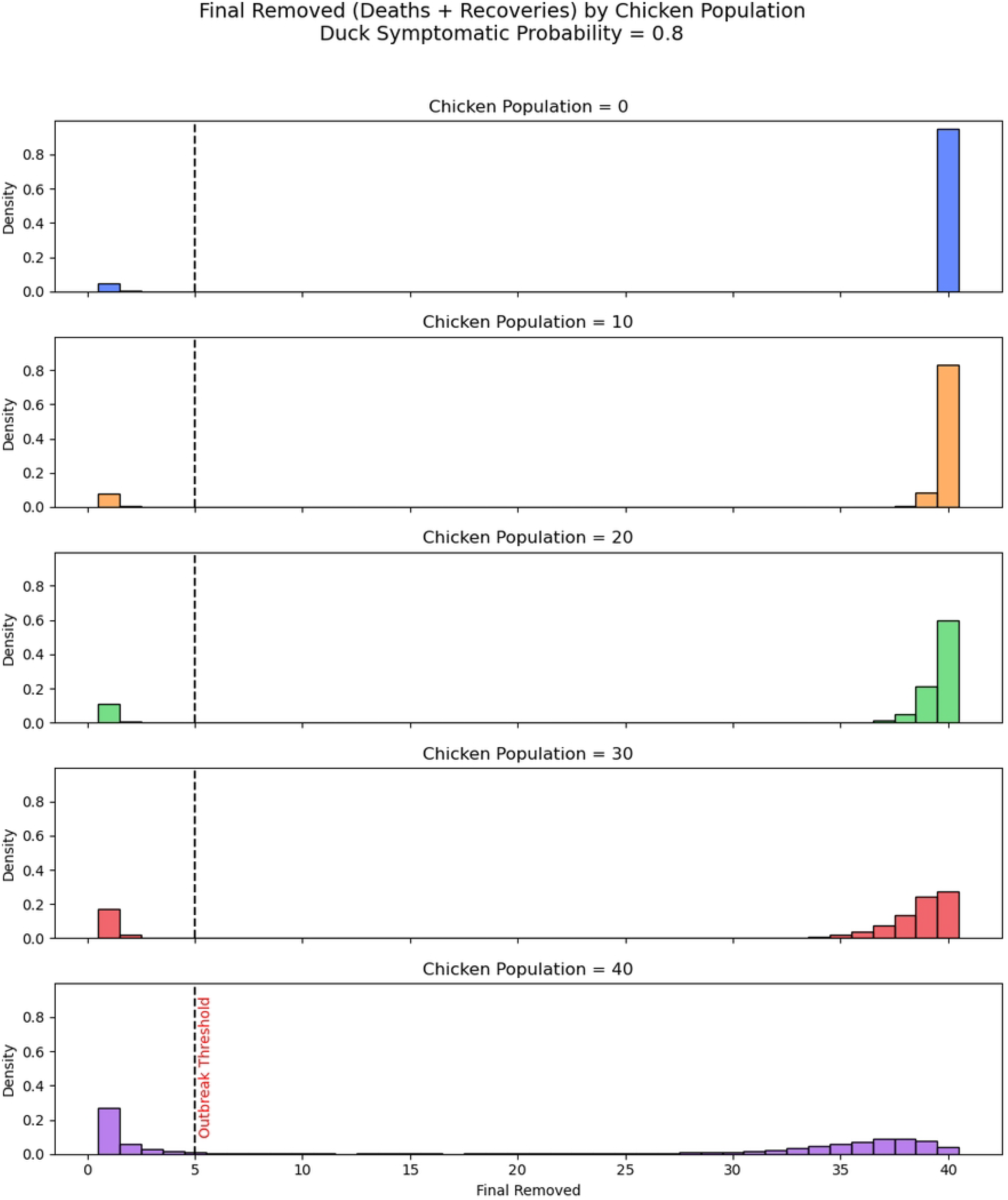
Histogram Final. *p*_*d*_ = 0.8 Histogram of distribution of final removed numbers (dead or recovered) with varied number of chickens in flocks of size 40. Duck symptomatic probability set to 0.8. The vertical dashed line indicates time series that are considered an outbreak (To the right of the dashed line).

**S5 Fig.**
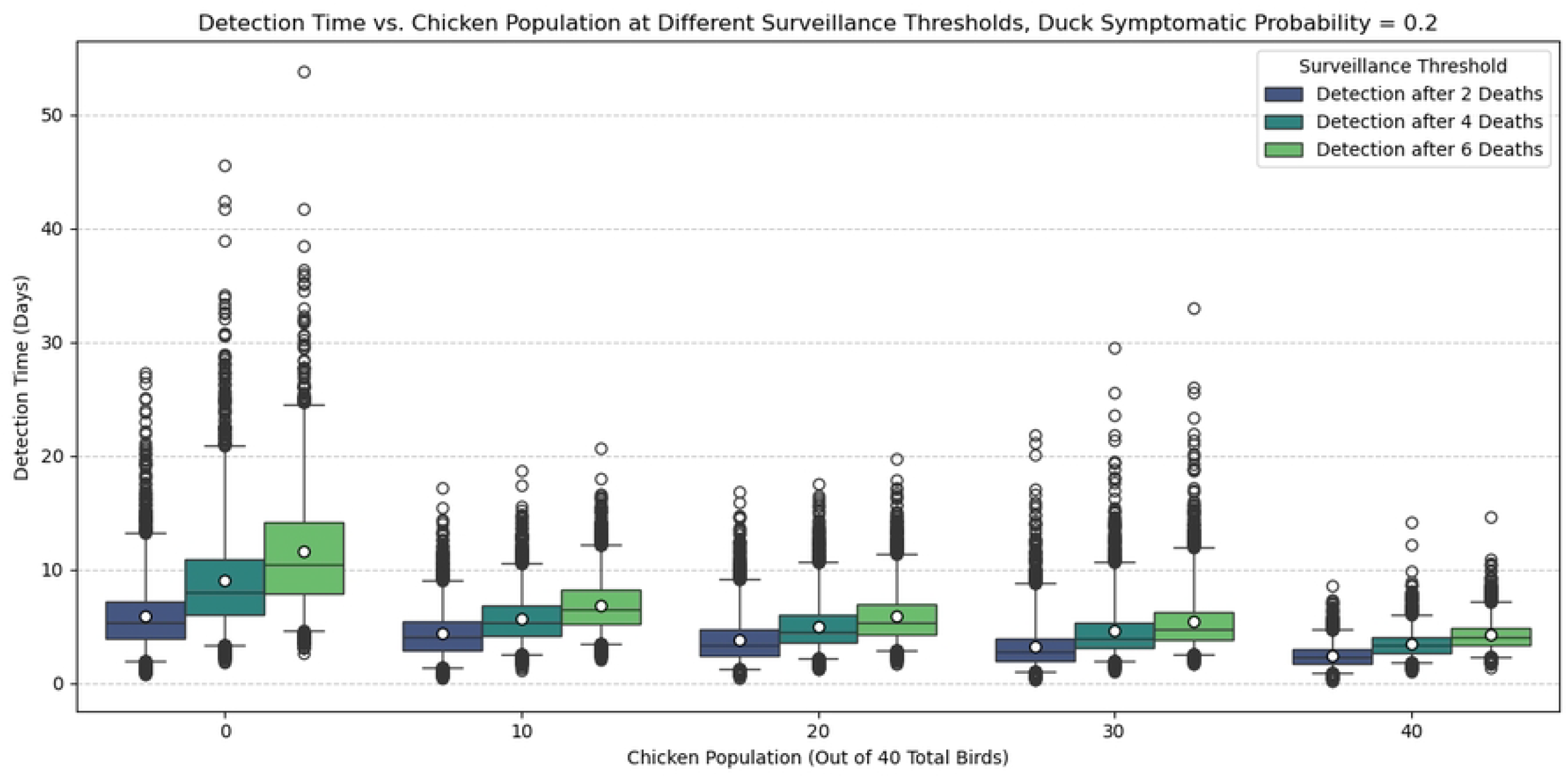
Box Plot Detection Time. *p*_*d*_ = 0.2, *κ* = 4 Detection time for different death thresholds for detection and chicken population out of 40 total birds, with *p*_*d*_ = 0.2. The boxes indicate the interquartile range (25th and 75th percentiles), and the whiskers indicate the 2.5th and 97.5th percentiles. This is similar to the box plots shown in the main manuscript, except with *κ* = 4 instead.

**S6 Fig.**
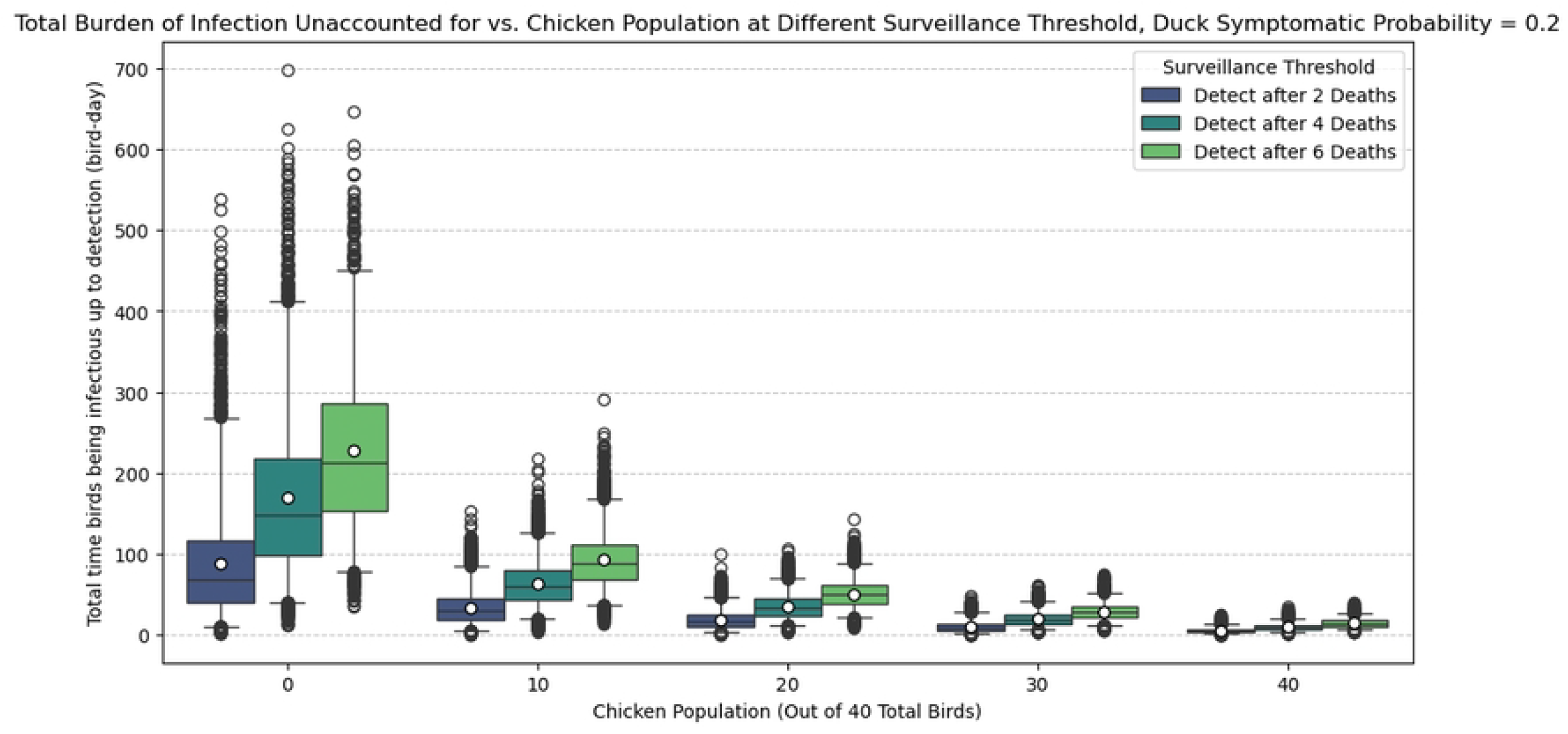
Box Plot Burden of Infection. *p*_*d*_ = 0.2, *κ* = 4 Undetected burden of infection (total time birds spent infectious up to the point of detection) for different death thresholds for detection and chicken population out of 40 total birds, with *p*_*d*_ = 0.2. The boxes indicate the interquartile range (25th and 75th percentiles), and the whiskers indicate the 2.5th and 97.5th percentiles. The white dot in the box indicates the average detection time. This is similar to the box plots shown in the main manuscript, except with *κ* = 4 instead.

**S7 Fig.**
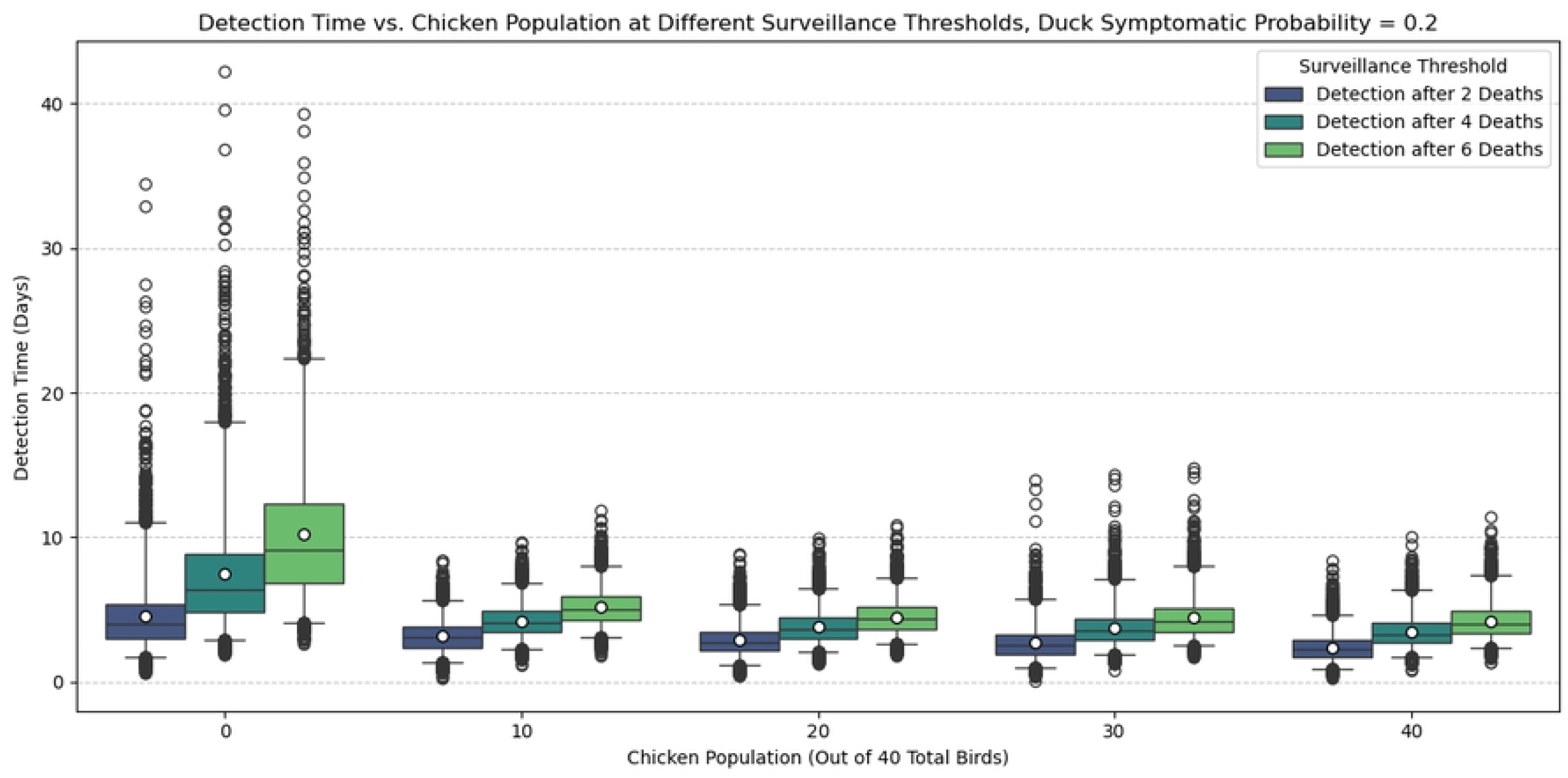
Box Plot Detection Time. *p*_*d*_ = 0.2, *κ* = 2 Detection time for different death thresholds for detection and chicken population out of 40 total birds, with *p*_*d*_ = 0.2. The boxes indicate the interquartile range (25th and 75th percentiles), and the whiskers indicate the 2.5th and 97.5th percentiles. This is similar to the box plots shown in the main manuscript, except with *κ* = 2 instead.

**S8 Fig.**
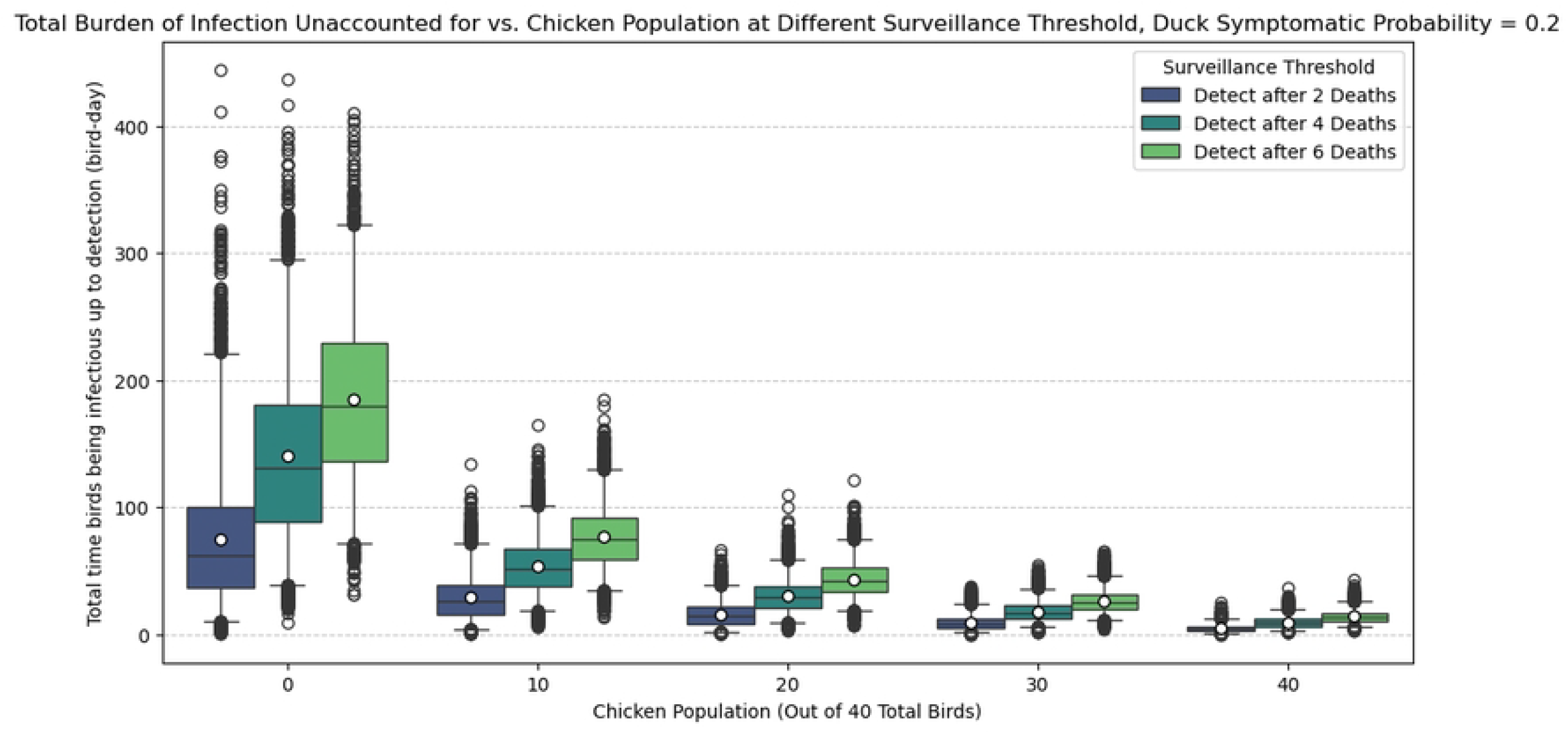
Box Plot Burden of Infection. *p*_*d*_ = 0.2, *κ* = 2 Undetected burden of infection (total time birds spent infectious up to the point of detection) for different death thresholds for detection and chicken population out of 40 total birds, with *p*_*d*_ = 0.2. The boxes indicate the interquartile range (25th and 75th percentiles), and the whiskers indicate the 2.5th and 97.5th percentiles. The white dot in the box indicates the average detection time. This is similar to the box plots shown in the main manuscript, except with *κ* = 2 instead.

## Notes

### Competing Interest Statement

The authors have declared no competing interest.

